# Mocap: Large-scale inference of transcription factor binding sites from chromatin accessibility

**DOI:** 10.1101/083998

**Authors:** Xi Chen, Bowen Yu, Nicholas Carriero, Claudio Silva, Richard Bonneau

**Affiliations:** Department of Biology, New York University; Department of Computer Science, New York University; Simons Center for Data Analysis, New York, NY 10010 USA

## Abstract

Differential binding of transcription factors (TFs) at *cis*-regulatory loci drives the differentiation and function of diverse cellular lineages. Understanding the regulatory interactions that underlie cell fate decisions requires characterizing TF binding sites (TFBS) across multiple cell types and conditions. Techniques, e.g. ChIP-Seq can reveal genome-wide patterns of TF binding, but typically requires laborious and costly experiments for each TF-cell-type (TFCT) condition of interest. Chromosomal accessibility assays can connect accessible chromatin in one cell type to many TFs through sequence motif mapping. Such methods, however, rarely take into account that the genomic context preferred by each factor differs from TF to TF, and from cell type to cell type. To address the differences in TF behaviors, we developed Mocap, a method that integrates chromatin accessibility, motif scores, TF footprints, CpG/GC content, evolutionary conservation and other factors in an ensemble of TFCT-specific classifiers. We show that integration of genomic features, such as CpG islands improves TFBS prediction in some TFCT. Further, we describe a method for mapping new TFCT, for which no ChIP-seq data exists, onto our ensemble of classifiers and show that our cross-sample TFBS prediction method outperforms several previously described methods.

## Introduction

A diverse host of regulatory factors bind to DNA to regulate gene expression and modulate the accessibility, functional status and structure of chromatin (1, 2). These factors form complex regulatory networks that underpin diverse patterns of cellular phenotypes (3, 4). Understanding the mechanistic basis of this regulatory, and ultimately phenotypic, diversity has remains a fundamental pursuit in biology and requires complete and accurate mapping of TF binding sites. Learning the patterns and processes of TF binding in a condition/cell type-specific manner is a critical step in identifying multi-factor targeted *cis*-regulatory modules (CRM) that are important in cell fate decisions (4, 5), and in understanding the causes and consequences of cellular rewiring in gene regulatory networks (6).

Most TFs are sequence specific (7) and TF sequence preferences are known (and readily available from multiple databases) in the form of position weight matrices (PWMs) (8, 9, 10, 11, 12). Today’s databases of PWMs cover a large number of factors spanning a multitude of species (8, 10, 11, 13). But TFBS identification based solely on PWM sequence matching is known for a number of problems (14). First, the length of a derived PWM is limited by experimental design, making it, in many cases, insufficient in describing the entirety of binding sequence environment. This is especially true in short probe-based technologies, such as protein binding microarrays, which often lead to the identification of incomplete sites or half sites (technology-dependent biases) (15). Shorter PWMs are invariably statistically more prone to false positive discoveries. Another concern is that degenerative sequences are known to be common among TFBS (16), but they are sometimes unaccounted for in classical PWMs, where terminal degenerative regions of a binding motif are removed. Additionally, motifs derived from *in vitro* high-throughput methods for binding site discovery do not capture patterns in the larger region surrounding motif sites that have recently been shown to be of importance for the bindings of a variety of TFs (17, 18). Lastly, high-throughput binding analyses carried out *in vitro* necessarily lack cell-type specificity, thus the binding profiles derived from such *in vitro* analyses do not reflect the dynamic chromatin landscape that promotes biologically meaningful binding events.

The lack of cell-type specificity in many TF binding assays is ameliorated by *in vivo* studies, such as ChIP-Seq. However, ChIP-Seq is carried out one TF and cell-type condition at a time, and its feasibility is often limited by factors such as the requirement to obtain high number of cells as input materials and the availability of high quality antibodies (19, 20). With the large number of factors and cell types, and the dynamic nature inherent in all TF binding events, it is challenging to capture the full scope of regulatory interactions for all factors and conditions with ChIP-Seq or any similar single-TF directed experiment.

Eukaryotic transcriptional regulation is shaped by chromatin dynamics, where accessible chromatin sets the stage for various types of regulatory interactions. Experiments that interrogate chromatin accessibility, such as digital genomic footprinting (DGF), DNase-Seq, ATAC-Seq and FAIRE-Seq have been used as promising alternatives to factor-specific ChIP-Seq for the identification of TFBS (21, 22, 23, 24). Because chromatin accessibility and nucleosome positioning are critical players enabling both the binding of TFs and the subsequent relay of regulatory information, such as co-factor recruitment and transcriptional machinery assembly, chromatin accessibility-based TFBS prediction methods has allowed cell type-specific predictions of binding sites for many TFs with a single experiment per cell type (25, 26, 27, 28, 29, 30). In spite of these advantages, the size and complexity of the mammalian genome, the diversity of TF behaviors (some TFs bind exclusively to nucleosome-free regions while others pioneer nucleosome-bound regions) and the large range of cell types (cell types modulate TF activity, TF-TF interactions and chromosome structure) make large-scale multi-cell type multi-TF binding site inference difficult, especially in a manner that balances method sensitivity and selectivity (31, 32, 33).

To address these challenges, we designed a TFBS prediction method that uses sequence-derived genomic features and one chromatin accessibility experiment per cell type to profile TFCT-specific binding activities. Our method has three components: 1) MocapG, an generic unsupervised method that ranks binding probabilities of accessible motif sites based on local chromatin accessibility, 2) MocapS, which integrates the motif-associated accessibility scores of MocapG with additional genomic features, such as TF footprints, CpG/GC content, evolutionary conservation and the proximity of TF motifs to transcription start sites (TSS) to train TFCT-specific predictive models under the supervision of ChIP-Seq data and 3) MocapX, which extends the selectivity of MocapS to more factors and cell types by mapping new TFCT conditions based on genomic feature distance to a nearest TFCT neighbor trained MocapS model using weighted least squares regression. The similarity-weighted ensemble prediction method, MocapX can connect TFCT-specific TFBS prediction models to TFCT pairs not directly queried using ChIP-seq or related methods. This cross-sample prediction framework, although limited to the scope of factors and cell types modeled, addresses the differences between TFCT conditions in TFBS prediction in a data-driven manner, and has the potential to expand the repertoire of putative TFBS with improved accuracy to any factors we have motif information for and in any cell type where chromatin accessibility data is obtainable. Additionally, we established a cross-assay comparison between model-based predictions using DNase-Seq and ATAC-Seq, in an effort to enable similar binding-site predictions from both of these widely adopted genomic technologies. In building a TFBS prediction method that learns and uses the differences between TF-chromatin interaction patterns, we hope to provide tools that help reveal the mechanistic complexity of mammalian gene regulation and chart the mammalian regulatory landscape spanning multi-lineage differentiation.

## Methods

### Obtaining candidate binding sites from motif collections

Human TF motifs (PWMs) were downloaded from the ENCODE motif collection (http://compbio.mit.edu/encode-motifs) and the CisBP motif database (http://cisbp.ccbr.utoronto.ca) (9, 10). We combined information from the two motif collections and filtered PWMs representing the same TF using pairwise comparisons based on normalized Euclidean distance (detailed in supplemental materials). The resulting nonredundant set of PWMs was then used to scan the human genome (hg19 assembly) to obtain candidate motif sites genome-wide using FIMO from the MEME Suite with options –max-strand –thresh 1e-3 (34). Overlapping motif sites (where at least half of a motif site overlaps with an adjacent motif of greater or equal length) are further cleaned to keep the motif site with a more significant matching score. Additionally, we excluded motif sites that overlap an ENCODE blacklisted region from downstream analyses (35).

### Chromatin accessibility and ChIP-Seq data processing

DGF and DNase-Seq data were downloaded from ENCODE as aligned reads (36). We filtered out reads with mapping quality lower than 25 and limited the number of mapped cuts per base pair to 50 to reduce the duplication effect caused by technical artifacts. DNase cut counts centered around each motif site were extracted from the processed BAM files with customized scripts (37), and all sites were cleaned with ENCODE mappability tracks prior to modeling to exclude unalignable regions of the genome from downstream analyses.

ATAC-Seq data for Gm12878 was obtained from Buenrostro et al. (23). We selected the experiment with the highest sequencing depth (SRR891268) to allow for sufficient sequencing depth for footprint detection. Reads were aligned using Bowtie with the option –best -X2000 -m1. As mentioned in (23), to extract the cut sites from ATAC-Seq BAM files, we offset the + strand read fragment by +4bp, and the - strand by -5bp. Similar mapping quality, mappability and 50 reads per base pair upper limit constraints were applied to the ATAC-Seq dataset.

ChIP-Seq broadpeak and narrowpeak tracks were downloaded from ENCODE. In case of multiple ChIP-Seq experiments for the same factor and cell type, tracks were merged to keep the intersections of all available experiments. Tracks flagged based on the quality metrics provided by the ENCODE consortium were excluded (https://genome.ucsc.edu/ENCODE/qualityMetrics.html) (38).

### The MocapG model

To obtain cell type-specific accessibility features associated with each motif site, we built a probabilistic model that classifies motif regions with a given number of cuts as either accessible or inaccessible. Briefly, we fit genome-wide accessibility cut count as a mixture of two negative binomial distributions and an additional zero component representing, respectively, accessible, inaccessible and zero-inflated regions of the chromatin.

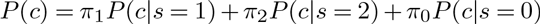

where

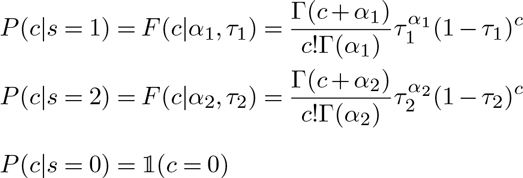

*c* is the number of DNase I cut count 100bp upstream and 100bp downstream of a specific motif site excluding the motif site itself. *α, τ* are the mean and dispersion parameters of negative binomial distributions respectively. π_0_, π_1_ and π_2_ correspond to the probability of the zero (*s*_0_), inaccessible (*s*_1_) and accessible (*s*_2_) component, where π_0_ + π_1_ *+* π_2_ = 1. Model parameters were estimated with an Expectation-Maximization (EM) algorithm, which takes a set of random initial parameter values (where *α*_1_ << *α*_2_ and π_1_ > π_2_ to avoid label switching) and genome-wide cut count in 200bp windows as input and outputs model parameters that maximize the log-likelihood function *log*(*P*(*c*)). Each motif region was then assigned a binary accessibility indicator *S* and a log likelihood score *L* based on the probability ratio of the region being accessible to inaccessible.

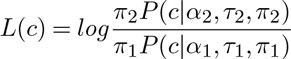

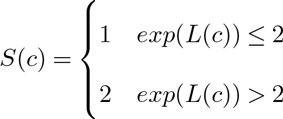

An *S*(*c*) of 2 corresponds to a likelihood ratio where a motif region is at least twice as likely to be accessible than inaccessible.

### Calculating other motif-associated genomic features

For all accessible motifs, we assessed the probability that a footprint profile exists around the motif site using a pair of binomial tests adapted from (28). For DNase-Seq, the binomial tests yield scores for strand-imbalance and depletion of DNase I cuts at motif sites. For ATAC-Seq, strand-imbalance is insignificant because of the absence of a size selection step. We merged strand information to assess if a motif region is depleted of cuts as compared to the left and right flanking region respectively. Detailed footprint score calculations are provided in the supplemental methods.

PhastCons and PhyloP scores were calculated for each motif region from conservation tracks downloaded from USCS (ftp://hgdownload.cse.ucsc.edu/goldenPath/hg19). TSS proximity features were calculated based on RefSeq Genes. Repeats were labeled for each motif region based on RepeatMasker calls (39). Mapability tracks were downloaded from ENCODE and selected based on corresponding DNase-Seq/ATAC-Seq sequencing length. Mapability scores, GC and CpG content were computed for a region 100bp upstream and downstream of each motif site. CpG islands were calculated as defined in (40, 41) (detailed in the supplemental methods). We provide customized scripts for all motif-associated feature calculations.

### Training sparse logistic regression classifiers to predict TFBS (MocapS)

We built sparse logistic regression models using the LIBLINEAR package in R to classify motif sites as true (overlapping with ChIP-Seq peaks) or false (not overlapping with ChIP-Seq peaks) binding site based on a list of genomic features and their interaction terms (42).

For all the motif-associated genomic features, we apply a correlation filter to retain, for each trained TFCT condition, 10 features that are most highly correlated with the ChIP-Seq signals based on the Pearson correlation coefficient (PCC). A second correlation filter was applied to the top 10 features and their second-order interaction terms to retain 30 most correlated features. Each TFCT condition was trained on a stratified sample of motif sites representing data from all except the hold-out chromosomes.

We adopt L1 regularization and tune the shrinkage parameter for each TFCT condition by performing 10-fold cross validation to optimize the area under the precision and recall (AUPR) curve. To avoid overfitting, the shrinkage parameter was further tuned such that the resulting classifier is sparser than an optimally performing classifier and still yields a near-optimum (within one standard error of the optimum) cross-validation performance. The final model parameters are estimated by aggregating over 100 bootstrap runs of such sparse logistic regression model fitting for each TFCT condition to reduce estimator bias. Model performance was assessed with data from the held-out chromosome. More details are provided in the supplemental methods.

### Cross-sample TFBS prediction (MocapX)

To expand binding site prediction to TFCT conditions where ChIP-Seq data is unavailable, we use robustly weighted least squares regression to derive mapping vector *β* to match new samples (samples without ChIP-Seq) to trained models for TFBS predictions. The regression problem is such that the derived sample weight 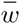 and mapping vector 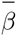 minimize the following error function

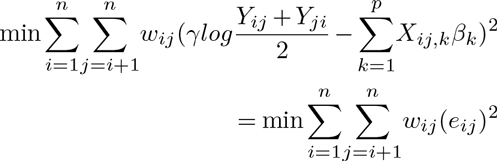

where *n* is the number of TFCT conditions; *p* is the number of features; *X*_*ij,k*_ is the binary probability that feature *k* from TFCT *i* and TFCT *j* are derived from the same distribution based on KS test for continuous features and Chi-square test for categorical features; *Y*_*ij*_ is the predictive performance (AUPR) of model trained in TFCT *i* when applied to classify motifs in TFCT *j*; γ is a hyperparameter that binarizes the target variable *Y* into sample pairs that cross-predict well and ones that cross-predict poorly; sample pair weight *w*_*ij*_ is fitted through iteratively re-weighted least squares regression using the rlm function in the MASS package in R with the Tukey’s bisquare family psi functions (43, 44).

We use 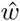 and 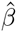 to compute a sample mapping *i* → *j* that maximizes 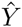 as below

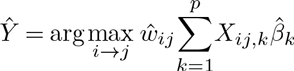

Mapping is then assigned if and only if 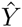 compares favorably with *Y*_*M ocapG*_ in TFCT *i*. More implementation details are provided in the supplemental methods.

### Comparison to other TFBS prediction methods

PIQ and CENTIPEDE were given the same motif input and accessibility data (25, 26). Because PIQ only makes predictions to sites that carry significant footprint profile, to allow a more complete comparison, we assign unpredicted sites the lowest posterior scores in PIQ. Comparisons are based on three metrics: AUPR, sensitivity at 1% false positive rate (FPR) and areas under the receiver-operating characteristic (AUROC) curve.

## Results

### Compiling a large unbiased set of candidate motif sites improves the sensitivity of TFBS prediction

The initial step of our computational pipeline involves the use of a large set of motifs from multiple sources to scan the genome to obtain TF-specific candidate binding sites. There is a many-to-many relationship between TFs and motifs representing them, as some motifs are derived using DNA-binding domains that are shared among several TFs (e.g. the motif for SP7 and SP9 are directly transferred from SP8 because of deep DNA-binding domain homology), whereas some TFs show diverged binding preferences that are matched by two or more distinct motifs (many TFs, e.g. NFKB, have a shared canonical motif accompanied by several non-canonical motifs) (10, 15, 45, 46). Among the 857 TFs represented by the motif collection, over half have three or more motif representations. The large number of motifs representing the same TF is resulted from either the intrinsic diversity in binding preferences or simply the differences in techniques used to derive them. For example, widely studied TFs, such as REST and CTCF, also tend to have large sets of motif representation, and consequently a large amount of redundancy between motifs (Figure S1B). To remove redundancy and resolve these many-to-many mappings prior to predicting binding sites, we introduced a similarity index based on normalized Euclidean distance. We compared the distance between vectorized motifs that represent the same TF and then kept the motifs from each similar cluster as described in the methods section (Figure S1).

We used a loose threshold (*P*-value < 1e-3) to scan for candidate binding sites. Given our overall scheme to subject identified binding sites to downstream classifications, a lower threshold for identifying putative sites can improve sensitivity with little increase in false positive rate. This lower threshold does incur a sizable computational cost, but we have found that this high level of inclusiveness is important for the following reasons. First, well-matched motifs only imply TF binding in a minority of cases. Non-concensus, low-affinity binding sites are sometimes required for proper functional readouts (47). We found for the majority of TFs, motif matching scores only weakly correlate with ChIP-Seq signal (average PCC is 0.05) (Fig. 3A). It was also shown through various studies for many different TFs that PWM matching score is a poor predictor of binding selectivity (25, 48). Thus using motif score to preselect for candidate binding sites will lead to sampling bias. Secondly, the number of motif occurrences or the occurrence of clustered motif sites are often important indicators of functional and tissue/cell type-specific binding. Many functional regulatory sites, especially in higher eukaryotes are found in clusters, either of themselves (homotypic clusters) or with each other (e.g. super enhancers) (49, 50, 51, 52). While site-wise degeneracy tends to be tolerated through evolutionary reshuffling, the formation of fusion sites and larger functional CRMs suggests that much of the complexity of mammalian TF binding lies, not in the qualitative matching of individual site, but in large part, in the arrangement of sites (53, 54, 55). Lastly, not all binding events require the physical interaction between TF and DNA molecule, thus a sequence-specific binding site may be absent at various sites when instead of binding directly and individually to the DNA, TF binds as a part of a protein complex or interacts with parts of the chromatin via chromatin-modifying enzymes (56, 57). For all of these reasons loosening the motif scan threshold can help achieve more complete coverage of binding sites on a genome scale (Figure S2) and subsequently allow more relevant genomic features, such as chromatin accessibility patterns, to drive the binding site prediction (Figure 3).

**Figure 1:**
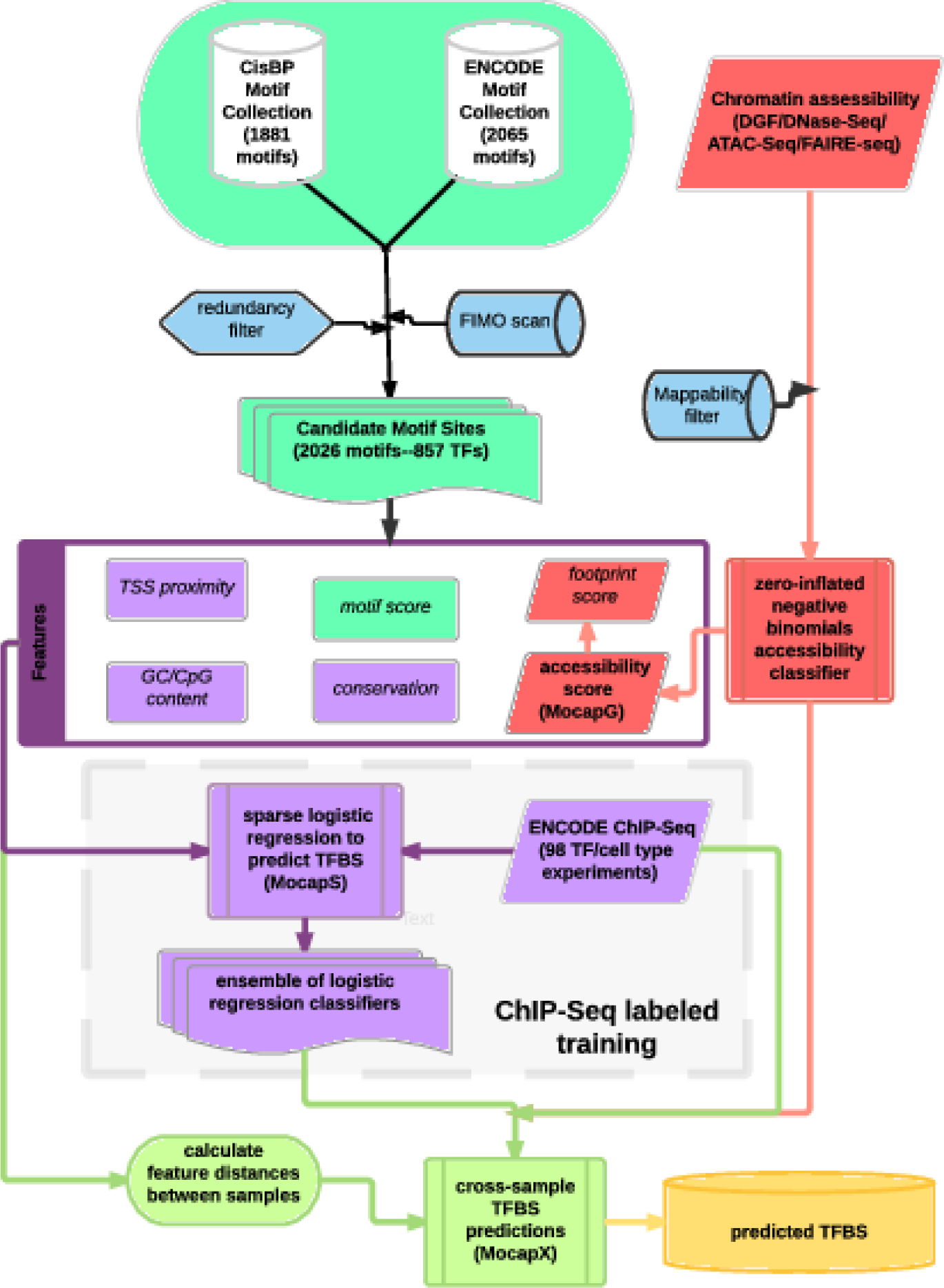
Our TFBS prediction pipeline. We compiled a non-redundant set of TF binding motifs, and compute genomic features for all candidate motif sites. We trained sparse logistic regression models to predict binding sites (MocapS) for 98 TFCT conditions, for which ChIP-Seq data is available in ENCODE cell type K562, A549 and Hepg2. True binding sites are defined as motif sites that overlap ChIP-Seq peaks. For a new TFCT condition, binding sites are inferred from either the unsupervised accessibility classifier (Mocap) or a trained sparse logistic regression classifier according to sample mapping using weighted least squares regression (MocapX). Shaded area stands for supervised training steps; unshaded area are steps for data acquisition (top) and making predictions (bottom).

**Figure 2:**
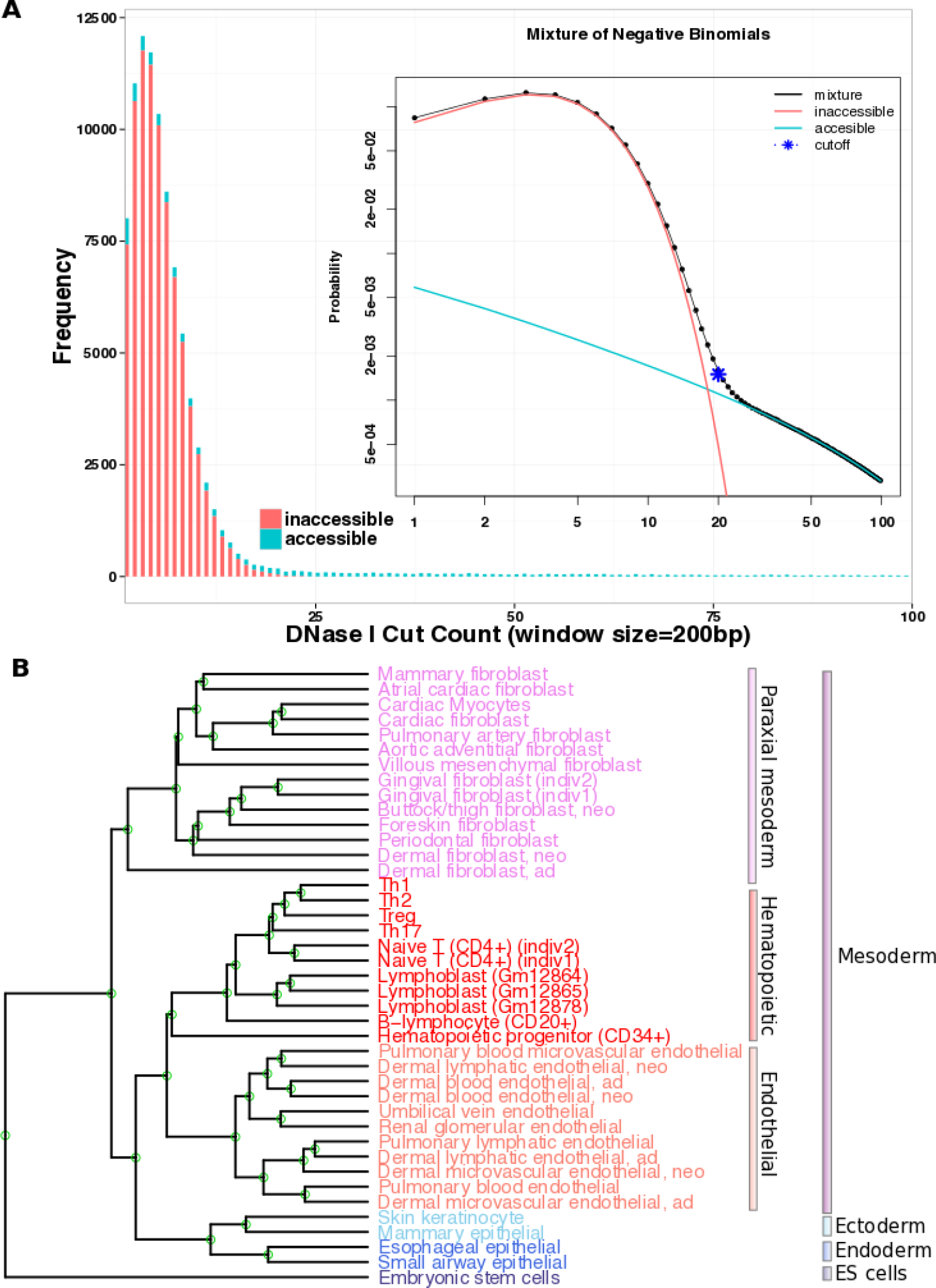
Modelling DNase I cut count as a mixture of negative binomial distributions. (**A**) Distribution of DNase I cut count simulated using zero-inflated negative binomial model parameters derived using an EM algorithm (n=100,000). Red: cut count from inaccessible regions of the chromatin; Blue: cut count from accessible regions of the chromatin. Inset: Cutoff point is determined by the probability ratio between accessible and inaccessible components. X and Y axes are in log scales. (**B**) Hierarchical clustering of the accessibility landscape of ENCODE cell types. Genome is binned into 400bp (overlapping by 200bp) windows, and the accessibility of each genomic window is classified using the zero-inflated negative binomial mixture model as 1s (accessible) and 0s (inaccessible). Cell types cluster in accordance with their developmental origins.

**Figure 3:**
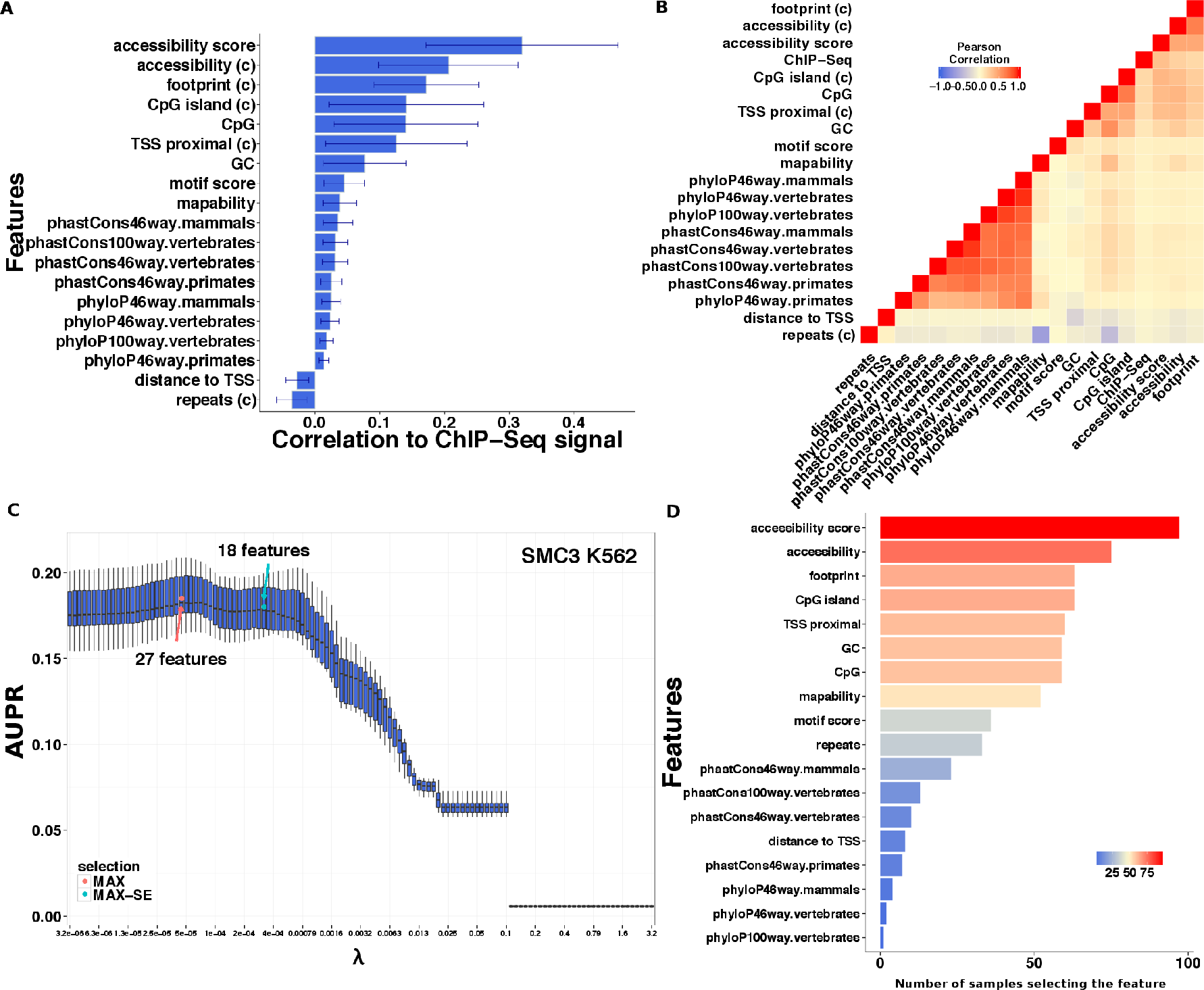
Feature selection and classifier training. (**A**) Genomic features ranked by their correlation to the ChIP-Seq signal. Barplot showing the average correlation for each genomic feature over 98 TFCT samples. Error bars mark average +/− one standard deviation. (**B**) Clustering heatmap showing PCC between genomic features across motif sites. Red: positive correlation, white: no correlation, blue: negative correlation. (**C**) Ten-fold cross-validation performance (AUPR) while adopting different shrinkage parameters λ. We tune the shrinkage parameter to approach maximum AUPR. Red dot marks the shrinkage level (sparsity) that corresponds to the maximum 10-fold cross-validation performance. Green dot corresponds to our selected feature combination–the sparsest model that achieved a near optimum (within one standard error of maximum) cross-validation performance. Example TF: SMC3, cell type: K562. (**D**) Barplot showing the number of times each feature is selected in the 98 trained models. Bar colors are scaled. Red and blue corresponding to more and less commonly selected features respectively.

### Classifying chromatin accessibility landscape with a mixture model of zero-inflated negative binomials recovers patterns of multilineage differentiation

Chromatin accessibility data are rich in cell type-specific regulatory information. To create a baseline method to rank motif sites based on chromatin accessibility (MocapG), we modeled chromatin accessibility states as a mixture of zero-inflated negative binomials, where two distributions for both accessible and inaccessible components of the chromatin are approximated with an EM algorithm. The negative binomial distribution was chosen because it models well the overdispersion commonly found in next-generation sequencing data (25, 58, 59), and has the flexibility to be applied to genomic regions of various sizes. This choice of distribution is also general enough to describe the signal-to-noise patterns associated with different experimental protocols for measuring accessibility (DGF, DNase-Seq, ATAC-Seq and FAIRE-Seq) (Figure S3). Further, we added a zero component into the mixture to assess the amount of zero inflation due to the lack of sequencing depth, e.g. those often found in FAIRE-Seq experiments. For a given accessibility experiment, we learned the component parameters from a cut count (for DGF, DNase-Seq or ATAC-Seq) or fragment count (for FAIRE-Seq) distribution randomly sampled from the mappable regions of the genome. The accessibility for a given genomic region (e.g. a motif region) is then decided based on the log likelihood ratio between the accessible and inaccessible components inferred (Figure 2A).

To test the mixture model’s ability to classify chromatin accessibility and distinguish between developmental cell types, we obtained ENCODE chromatin accessibility data (DGF or DNase-Seq) for 41 normal human cell types at various stages of development. For each cell type, we binned the genome into 400bp windows, and used our model to classify regions of the genome into a binary profile of ones (accessible) and zeros (inaccessible). We then calculated pair-wise Euclidean distance between cell types and performed hierarchical clustering on the resulted accessibility profile. As shown in Figure 2B, cell types fall into clusters based on germ layer origin, confirming the utility of the classification method across experiments with various sequencing depth. The result also exemplifies the key role chromatin accessibility plays in coordinating cellular differentiation and directing cell fate decisions (60). MocapG was then used to generate our main motif-associated accessibility features in logistic regression training.

### TF footprint profiles are condition-dependent

Because of the overarching role of chromatin accessibility in directing TF regulation and distinguishing cell type identity, we next sought to model footprint patterns surrounding accessible motifs to determine if there is evidence of direct physical binding of a TF to each candidate motif site. Using a pair of binomial tests, we assessed whether a footprint profile exists (if the motif site is depleted of cut count as compared to its left and right flanking regions respectively). To examine the strand-specific patterns of footprint scores across TFCT conditions, we plotted the averaged cut count profile surrounding motifs found in ChIP-Seq peaks (Figure S6, Appendix). Due to the occlusion of the immediate flanking nucleosomes, the 5’ cut distribution at binding sites tend to be strand-biased, with the positive-strand cuts more likely to accumulate on the left flanking region of a site, while negative-strand cuts more on the right (28). The fragment size selection step in DGF protocol further accentuate this strand imbalance towards the center of binding sites when compared to ATAC-Seq (size selection step is absent from a typical ATAC-seq protocol) (21, 23). We thus adopted a strand-specific footprint detection method for DNase I footprints detection and a non-strand-specific method for ATAC-Seq footprint detection (see Methods).

Previous work has pointed out limitations in footprint modeling, showing that the footprint signature is subject to several systematic biases including enzyme cutting and sequence bias (30, 61, 62, 63, 64, 65). Corroborating these observations, we found that DNase I cuts tend to result in more conspicuous footprint than Tn5 transposon insertion cuts, as shown in the differences in cut count profiles of factors, such as CTCF and RAD21 (Figure S6A-B). As the behavior of these two factors does not vary significantly among cell types measured by DGF (K562, A549 and Hepg2), the observed differences between DGF-measured and ATAC-Seq-measured cell types are likely *bona fide* differences between the enzymatic activities of DNase I used in DGF and Tn5 transposase used in ATAC-Seq (61).

Aside from technical complications, we observed high variability in the shapes and patterns of DNase I footprints across factors and cell types. Some factor-specific signatures are attributed to the sequence properties of the motif and/or the binding behavior of the TF. For example, FOXA1, FOXA2, MXI1, CEBPB, NFYA and RFX5 show DNase I footprints that are center-inflated rather than exhibiting a canonical center-depletion (Figure S6, Appendix). These factors tend to have A/T rich motifs and three out of the six TFs (FOXA1, FOXA2, NFYA) have been previously implicated as pioneer factors (26, 66, 67, 68, 69). These inflated footprints might suggest transient binding dynamics, due to the competition between destabilized nucleosome and TFs, or between interacting TFs. Some TFs, e.g. ZNF384 and ARID3A have highly noisy cut patterns surrounding motifs. This transient binding might be due to either poor motif/ChIP-Seq quality or the absence of consensus sequence preference, as these motifs also correspond to the samples with incomplete ChIP-Seq coverage (Figure S2). Differences in footprint profiles can also result from cell type-specific TF activities, e.g. due to the presence or absence of a co-binder. Because ChIP-Seq cannot distinguish direct from indirect binding, a TFCT condition lacking an average footprint profile might suggest the dominating effect of indirect binding or low affinity binding of the TF in the tested cell type. In addition, the lack of an obvious footprint could also be due to the fact that our motif-centric method, by definition, requires footprint discovery be anchored around the center of a known motif, which might not apply to all TF motifs. This is especially true for clustered binding sites where the occurrence of multiple motifs (10-30bp) within one ChIP-Seq peak region (around 200bp) makes it difficult to pinpoint with either motif matching, accessibility level or footprint profiling which narrow region corresponds to the actual physical binding of the TF (70). Although centering footprint detection around motifs can absorb part of the variation seen across factors and cell types, the diversity of TF-specific footprint profiles necessitates developing more flexible and condition-specific footprint detection methods, especially for ATAC-Seq footprints.

### Building TFCT-specific binding site models

To build condition-specific TFBS prediction models, we selected 19 motif-associated genomic features and trained sparse logistic regression classifiers (MocapS models) under the supervision of ChIP-Seq for 98 TFCTs (representing 52 TFs). Across the samples, local chromatin accessibility scores are the most consistently correlated with ChIP-Seq signal (Figure S4) and also the most consistently chosen by our model selection procedure. Here we defined local chromatin accessibility as a relatively small region 100bp upstream and 100bp downstream of a motif site, excluding the motif site itself, as we found chromatin accessibility tends to taper off after 100bp and the motif site itself tends to be depleted of DNase I cuts because of TF footprint (Figure S6). In comparison to local chromatin accessibility scores, our binary accessibility feature based on the negative binomial mixture model and footprint scores correlate to ChIP-Seq in a less consistent manner (Figure S4).

GC/CpG content-associated features, namely, GC content, CpG count and CpG islands also exhibit more divergence among TFCTs, with TFs, such as E2F6 and HEY1 showing ChIP-Seq correlation to GC/CpG content comparable with accessibility features, while GC/CpG content around MAFK and MAFF motifs are barely correlated with ChIP-Seq (Figure 3A-B,S4). Additionally, a binary feature separating motif site based on TSS proximity (1kb) tends to be more correlated with ChIP-Seq signal than the continuous feature distance to TSS (Figure 3A-B,S4). This suggests a non-linear relationship exists between the distance of *cis*-element to a nearest TSS and its regulatory activity, and that for some TFCT conditions, a disproportional majority of regulatory interactions take place in TSS proximal sites. Evolutionary conservation scores only weakly correlate with ChIP-Seq signal, despite the common assumption that functional *cis*-regulatory elements are more likely to be conserved (Figure 3A-B and Figure S4). Among the two types of conservation scores and three different evolutionary distances we tested, phastCons scores, which measure the probability of a motif site belonging to a conserved element seemed to be more predictive of TFBS than phyloP scores that attempt to capture accelerated rate of evolution. Compared to vertebrate and primate conservation scores, mammalian evolutionary conservation scores were found to be the most correlated with and predictive of TF binding (Figure 3A,D). Mapability scores and repetitive sequences both appeared to correlate with CpG features and accessibility and likely confound binding site predictions (Figure 3B).

To incorporate relevant features in binding site prediction and preclude spurious signals, we filtered genomic features based on their correlation to ChIP-Seq signal (detailed in methods) before subjecting them to the sparsity constraints in logistic regression (Figure 3A). Further, we included interaction terms, reasoning that less directly correlated features, such as TSS proximity and evolutionary conservation scores could have modulating effects on TF binding. We used L1-regularization to constrain model sparsity, selecting models that achieve good cross-validation AUPR but also have the potential to generalize well out-of-sample (Figure 3C). For the 98 samples we trained on, there is a general agreement between a feature’s correlation with target variable (ChIP-Seq) and the likelihood of the feature being selected into the final model (Figure 3A,D). Among the individual and interaction features, we found features involving local chromatin accessibility to be the most widely selected predictor of TF binding across TFCT conditions (Figure 3D,S5).

### GC/CpG content surrounding TF motifs modulates TF binding

Sequence features affect TF binding through mechanisms such as altering local DNA shape, affecting nucleosome positioning, DNA methylation and/or influencing co-binding sites (17, 71, 72, 73). In contrast to the good generalization properties of chromatin accessibility features, sequence features tend to influence TF binding in a TFCT-dependent manner (17). Among all possible k-mers, the most prominent sequence features with predictive power are GC content, CpG dinucleotide frequency and stretches of CpG islands. We reason that the incorporation of key features as such can provide quantitative descriptions of the local sequence/chromatin environment. For example, GC/CpG content can act as a proxy for nucleosome occupancy and DNA methylation, both of which are known regulators of TF binding (74, 75, 76).

As we observed high between-sample variation in correlations between GC/CpG features and ChIP-Seq signal (Figure S4), GC/CpG feature usage varied across factors and cell types (Figure S5C). Among the 98 trained TFCT experiments, we found that GC/CpG sequence features emerge, most frequently, as an interaction term with accessibility features (Figure S5B). Given the dominant and relatively universal usage of accessibility features, GC/CpG sequence features seemed to play a modulating role boosting the predictive performance of chromatin accessibility for some TFCT conditions. For example, E2F family factors appeared to prefer CpG rich sequence environment almost universally, whereas the Forkhead family factors, such as FOXA1 and FOXA2, tend to make use of the GC/CpG features in a less straightforward manner.

Further, we saw a moderate correlation between GC/CpG sequence features and TSS proximity (Figure 3B), as well as a significant overlap between the usage of these two types of features across factors and cell types in our trained models (Figure S5C). This is consistent with the fact that most promoter regions are GC-rich. In FOXA2 and ZBTB7A, for example, the effect of GC content on nucleosome occupancy and TF binding appears to depend on TSS proximity, with TF binding at TSS proximal sites featuring a more positively correlated relationship with GC/CpG content than TSS distal sites (positive coefficient for interaction features between GC/CpG and TSS proximity and negative coefficient for GC/CpG feature). This is reminiscent of what is observed in macrophage pioneer factor PU.1, where high GC content promotes the stable positioning of nucleosomes and leads to greater nucleosome occupancy at PU.1 sites, but very high GC content (e.g. CpG islands) disfavors nucleosome assembly at proximal sites and low GC content at distal sites (77).

Overall, our trained models show diverse usage of GC and CpG sequence features across factors and cell types (Figure S5). Because GC content surrounding motif sites was shown to agree with core motif GC content in a TF family-specific manner, the sequence feature preference could be an indirect result of a homotypic environment for binding and cooperativity (17). Also, extremely GC-poor regions are thermodynamically disfavored for nucleosome positioning due to the stiff property of poly(dA:dT) sequences, so the wide-spread usage of GC content feature could underlie a structural basis for TF binding (78).

### Cross-sample TFBS predictions

To extend the usage of our TFCT-specific logistic regression classifiers to new factors and cell types, we next tested whether we can apply our trained models to the binding site predictions of TFCT conditions in the absence of ChIP-Seq data. We first evaluated the extent to which TFCT condition spesific models can cross-predict each other, where cross-prediction is quantified by measuring how well we can use model trained in TF *i* to predict binding for candidate motifs of TF *j*, and vice versa. To reveal the factor-specific contribution to cross-sample TFBS prediction, we collapsed data for the same factor across cell types and generated a distance matrix between prediction performance across the 52 TFs (Figure 4). Among the TFs we have trained on, factors can be broken into three major clusters based on cross-sample predictive performances (these clusters contain TFs that are well predicted by similar sets of genomic, motif and accessibility features). The largest cluster (red) has accessibility and GC/CpG sequence feature as the most prominent predictors. Cluster 2 (green) features the cohesion complex factors, CTCF, RAD21 and SMC3 together with MAF and USF family factors that all tend to make significant use of motif scores. Cluster 3 (blue) is comprised predominately of enhancer-associated regulators, such as EP300 and TEAD4 whose binding sites are more likely to extend beyond TSS vicinity, and pioneer factors, such as AP1 (JUN/JUND) and Forkhead family factors. We found that, despite the sparsity constraint, more complex multi-featured models are preferred by factors in this cluster.

**Figure 4:**
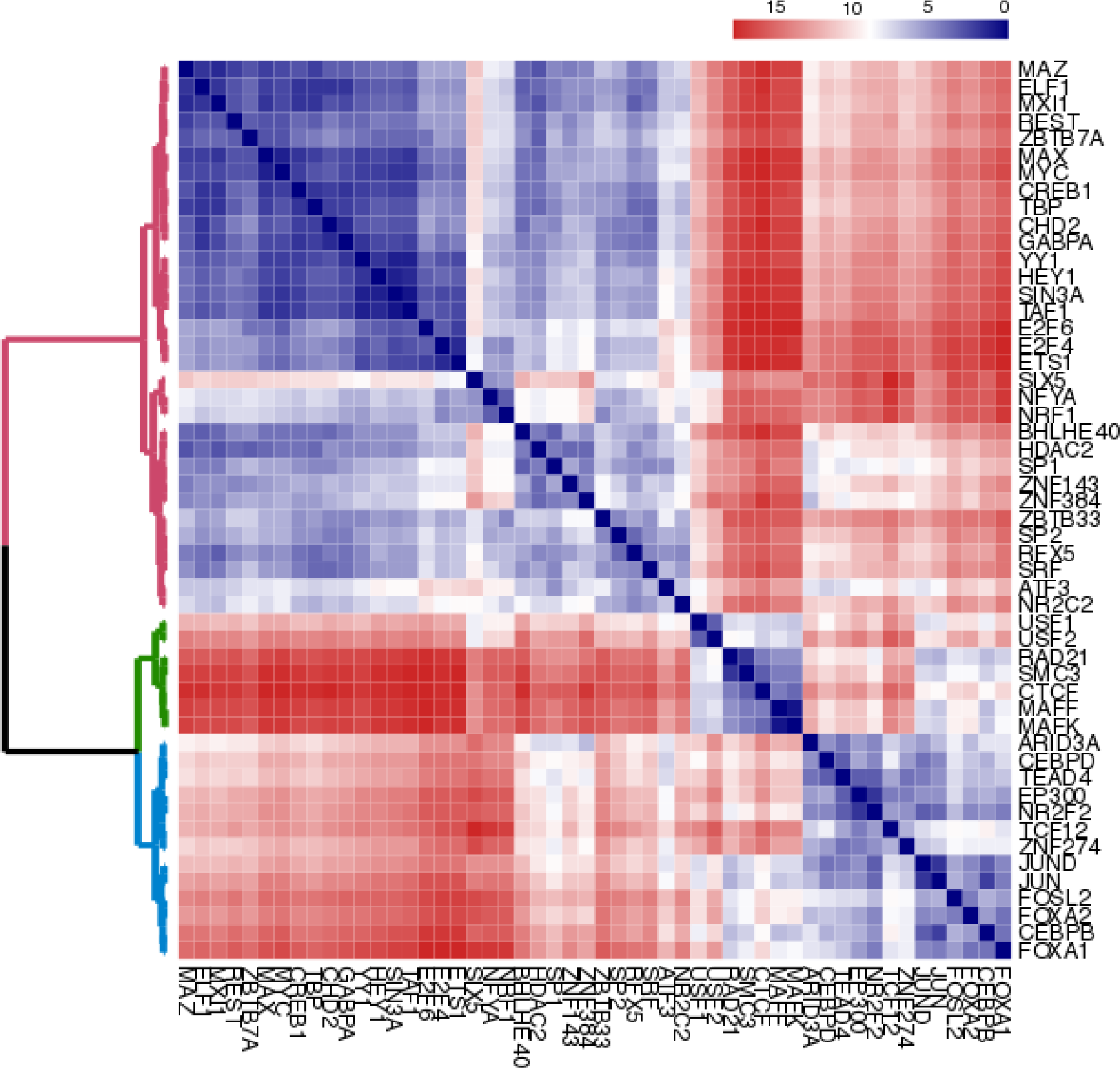
Heatmap clustering TFs based on the Euclidean distance between cross-TF prediction performances (AUPR). Red indicates large Euclidean distance and relatively poor cross-prediction performance between TFs; Blue indicates smaller Euclidean distance and good cross-prediction performance (where cross-prediction is the use of TF’s MocapS model to predict another TF’s binding). TFs are clustered together if they are more likely to share the same sparse logistic regression models for predicting TFBS. Data from multiple cell types, if available are averaged out for each TF.

It is worth noting that part of the cross-sample prediction performance trade-off stems from differing signal-to-noise ratios between samples. There is a high level of cell type-specificity even for the same TFs due to varying levels of TF activities and multi-factored interactions. We found that models trained in cell types where the TF show more true binding sites (ChIP-Seq peaks) tend to be more robust when applied to the same factor in another cell type. This is because TFs with fewer binding sites across the genome tend to produce more biased models. SIX5, for instance, has a very small set of binding sites in K562. This small true class of binding sites, although weight balanced with the false class, can only give us partial information about the factor’s binding preference, so it is often difficult to infer generalizable models or perform fair model evaluation on such datasets. To circumvent this bias and down-weigh such outlier samples, we used iteratively weighted least squares regression to derive a feature mapping vector (Figure 5). As the system is overdetermined (n ≫ p), we reason that down-weighing noisy outliers (samples with large residuals) can produce more robust sample mapping. Additionally, because not all supervised MocapS models show significantly improved performance over unsupervised Mocap, we constrained the use of this weighted mapping (MocapX) to models that showed significant improvement over unsupervised MocapG to control for uninformative mappings.

**Figure 5:**
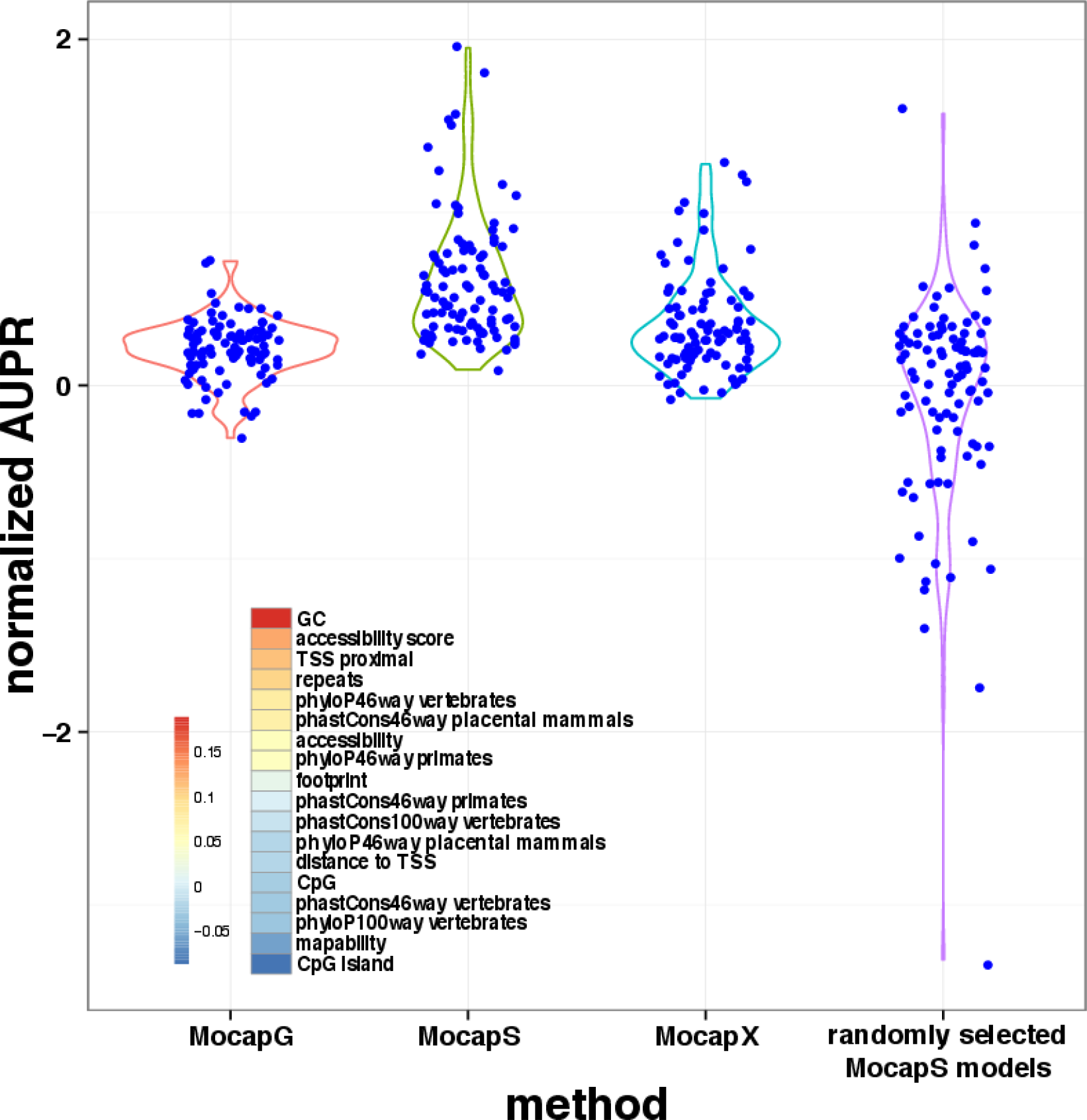
Cross-sample binding site prediction. Violin plot showing the hold-one-out performance for MocapX in comparison to MocapS (with MocapS models trained in the TFCT), MocapG (with local chromatin accessibility feature only) and randomly selected MocapS model (with random mappings between leave-out TFCT and MocapS model ensemble) performance. Inset: Heatmap showing weighted feature vectors that is used to compute distances between new TFCTs and TFCTs for which MocapS models have been trained. If no fit model exists in the trained model pool (no model is predicted to outperform unsupervised MocapG), MocapX will use MocapG for TFBS prediction.

The weight distribution in MocapX derived feature vector suggests cross-sample mapping is predominately driven by GC/CpG sequence feature similarities (Figure 5). Among the 98 leave-one-out mapping experiments, we were able to map 35 samples to another TFCT trained model (the rest chose the generic method MocapG). Fourteen of the 35 mappings were between cell types of the same TF, the rest mapped cross-TF. Among the cross-TF mappings, the most prominent one is between CTCF and RAD21 (Table S1). This is consistent with the overlapping roles of these two factors in defining chromosomal domains and in interactions with other cohesion complex components, such as SMC3 (79, 80, 81). It was shown that CTCF and RAD21 bind to a subset of their accessible motifs and a small number of these binding sites are further influenced by cell type-specific DNA-methylation (73, 82). MocapG was found to be rather inaccurate when predicting the binding of these factors. Incorporating motif scores and footprint profiles significantly improves prediction accuracy in these cases. Other examples of cross-TF mappings are ETS1/YY1, CEBPB/JUN, REST/SIN3A and MAX/MYC (Table S1). These factors were previously recognized as binding partners, so their binding sites likely co-cluster and thus share similar sequence environments and model preferences (83, 84, 85, 86). As a point of comparison, Figure 5 also demonstrates the peril of wrongly assigning models to samples: randomly selecting MocapS models for cross-sample predictions can negatively impact performance. This underlies the importance of discriminating between condition-specific models. Further, we found that inferring binding for TFs in conditions/cell types where they are inactive, partially active or have altered binding activities tend to confound cross-sample TFBS predictions (Table S1 and Figure S7B), potentially highlighting the need to integrate alternate data-types, such as TF expression or approaches that can explicitly account for TF activity in different cell types (87, 88). Taken together, we show that MocapX, although limited by the number and diversity of TFCT-specific models we have trained, presents a novel framework to generalize trained sparse logistic regression models to an increasing number of TFCTs with improved accuracy.

### Performance comparisons

We compared the performance of Mocap with two other motif and chromatin accessibility-based TFBS prediction methods CENTIPEDE and PIQ. As motif datasets show significant sample class imbalance, with true binding sites taking up only a fraction of the total candidate sites in our predictions (positive/negative ratio < 0.01%), AUROC scores are likely biased towards assessing correct classification of the majority class (which in our case is the non-binding sites that we are less interested in). Thus, we computed the AUPR for each leave-out TFCT to compare the predictive accuracy among methods (trade-off between precision and recall). We also evaluated sensitivity at 1% FPR to evaluate binding site prediction at a low false positive rate cutoff (trade-off between sensitivity and specificity) (Figure 6A and Table S1).

**Figure 6:**
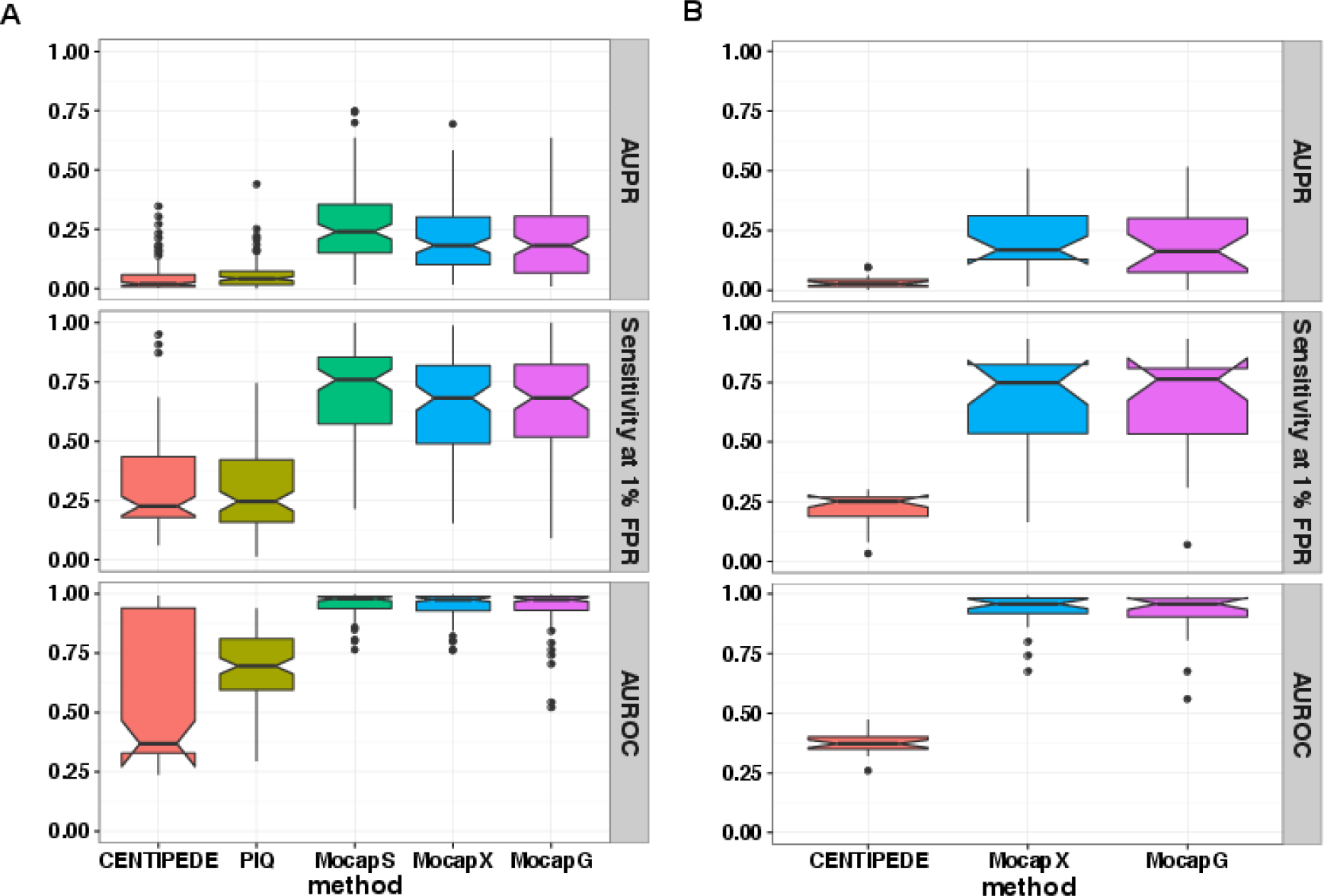
Method comparison between Mocap, CENTIPEDE and PIQ (98 TFCT samples in hold-out chromosome 15). (**A**) Boxplot showing overall performance of CENTIPEDE, PIQ, MocapS, MocapX, and MocapG method in predicting TFBS (n=98). (**B**) Boxplot showing performance of Mocap and CENTIPEDE applied to ATAC-Seq data in Gm12878 (n=23). Performance metrics used are AUPR (top panel), Sensitivity at 1% FPR (middle panel) and AUROC (bottom panel).

**Figure 7:**
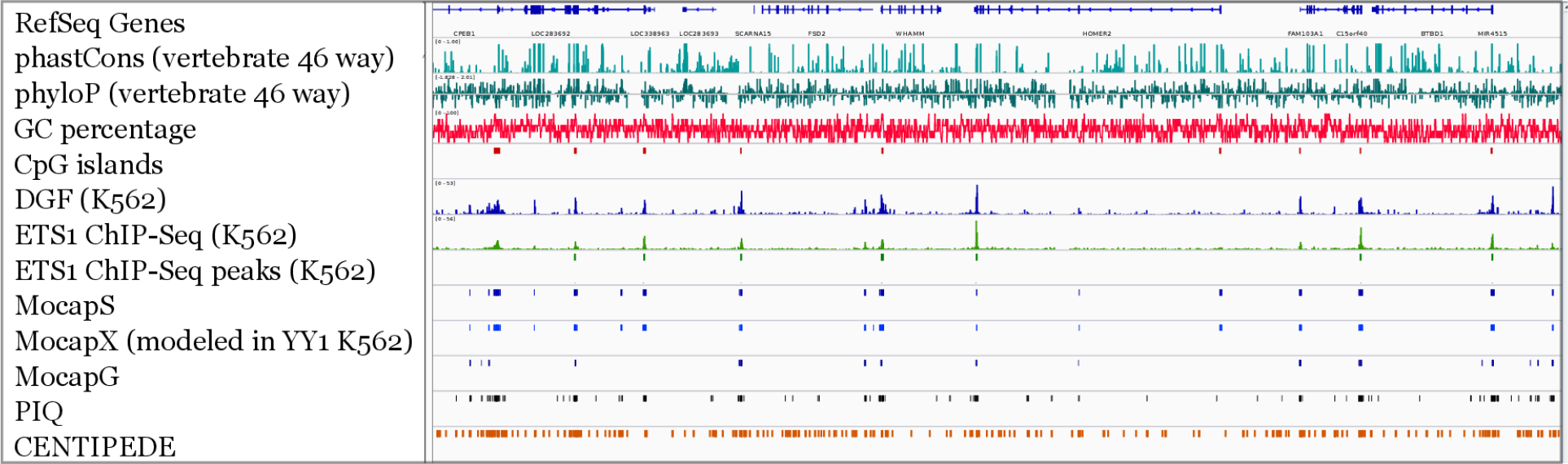
Genome browser view of predictions made by different methods. Tracks highlight region 85081291-85557900 on chromosome 15 for binding site predictions of ETS1 in K562. We standardized MocapG, MocapS and MocapX (modeled with YY1 in K562) prediction scores into z scores and used a cutoff of z > 3. Cutoff for PIQ and CENTIPEDE are 700 and 0.99 as suggested.

When compared to CENTIPEDE and PIQ, both our sparse logistic regression training-based MocapS and our extended MocapX methods showed a good balance between sensitivity and specificity across factors and cell types (Figure S7A-B). MocapG, which uses simple accessibility cut count ranking, although lacking in the completeness of coverage, remained a robust prediction method across factors and cell types (Figure 6A). But, MocapG did tend to fall short in TFs such as CEBPB, E2F4, RAD21 and MAFF where motif matching scores, sequence features or TSS proximity played major roles (Table S1). These cases clearly demonstrate the need to build TFCT-tailored models that use a diverse collection of features.

CENTIPEDE and PIQ both model footprints in great detail. PIQ uses refined technique to model motif and DNase I footprint. CENTIPEDE integrates motif scores, TSS proximity and evolutionary conservation with DNase I footprint in a hierarchical model. Overall, both methods rely heavily on the modeling of footprint profiles. So, when we apply these methods to TFBS prediction with a loosened constraint on motif matching score to improve sensitivity, they often fell short in ranking precision (poor AUPR scores). This points us to the limitation of methods that rely solely or heavily on footprint profiling. First, it is difficult to profile TF footprints even with the aid of factor-specific motifs, especially in a manner that balances specificity with completeness of prediction (due, as we discuss above, to a host of other influences like chromatin context and cell-type specific co-factors). Footprint detection methods, although descriptively useful, tend to be highly biased and condition-specific, thus lack the ability to generalize across factors and cell types and are often unfit for application on a global scale, to facilitate efforts, such as genome-wide CRM detection or gene regulatory network inference. Secondly, part of the variation in footprint profile might be intrinsic to differential TF binding activities. Although condition/protocol-specific bias correction using naked DNA could help resolve footprint patterns, the condition specificity of TF footprints both within and across TFCTs will likely remain prominent (27, 30). This again signifies the need for building predictive models that specifically address the differences between TFCTs.

### Adapting Mocap to predict TFBS using ATAC-Seq

ATAC-Seq has emerged in recent years as an efficient method to assay chromatin accessibility and typically requires much smaller cell sample than DNase-Seq to achieve comparable sequencing depth. This has led to the wide application of ATAC-Seq to clinically relevant rare cell types and conditions that were previously inaccessible to DNase-Seq due to limited cell numbers. The higher sensitivity and stable nature of Tn5 also results in more ambiguous/arbitrary footprint profiles in comparison to DNase I (Figure S6, Appendix).

To examine performance across assays, we applied MocapG and MocapX to ATAC-Seq data in human Gm12878 cells and compared the performance of Mocap with that of CENTIPEDE (Figure 6B and Table S2). MocapG, because of its simplicity, performs comparably across assays, whereas the differences in footprint profiles limit the specificity of MocapX when applied to ATAC-Seq. This loss in predictive performance is more pronounced for CENTIPEDE, which was designed specifically for DNase footprints. Despite the differences in footprint profiles, MocapX was shown to improve the performance of TFBS predictions for TFs, such as CTCF, RAD21 and USF1, where MocapG tends to perform poorly. This represents, to our knowledge, the first TFCT-specific effort to use ATAC-Seq for TFBS prediction.

## Discussion

In this work, we developed a DNA binding motif and chromatin accessibility-based method to predict cell type-specific TF binding. In designing a generalizable TFBS prediction method, we followed several key design principles: 1) we address the differences in binding behaviors between TF-cell type conditions, 2) we predict TFBS in a manner that balances method precision with recall, and 3) we employ approaches that improve method scalability. To distinguish between TFs and cell types, we incorporated in our analyses a range of motif-associated genomic features, including motif matching scores, chromatin accessibility, TF footprints, GC/CpG content, TSS proximity and evolutionary conservation. We assessed each feature’s contribution to TFBS prediction, and applied model selection to the training of an ensemble of TFCT-specific classifiers integrating these genomic features. We show that incorporation of sequence features, such as GC/CpG content surrounding TF binding motifs, significantly improves predictive performance and helps identify TFCT conditions sharing similar predictive models. To improve the sensitivity of our predictive method (recall), we start with a more complete coverage of binding sites by loosening motif matching threshold; and to provide more informative ranking of our predicted binding sites (precision), we set the objective function in sparse logistic regression training to optimize AUPR.

Our TFCT-specific models perform favorably in binding site predictions for a range of TFs and in a multitude of cell types in comparison to several previous methods. We show that this specificity and performance can be extended to other TFs that lack ChIP-seq via cross-sample TFBS prediction (MocapX) using condition-specific models trained in other TFCTs. Lastly, to promote scalability, we focused on designing a method that is computationally efficient and relies on only a single type of genomic assay, either DNase-Seq or ATAC-Seq to capture the chromatin dynamics around binding sites. Although a wide variety of functional genomic assays, such as histone modification ChIP-Seq, MNase-Seq and bisulfite sequencing could all contribute to TFBS prediction (89, 90), we chose to limit our required inputs to chromatin accessibility experiments, because they remain, to our knowledge, the most generalizable genomic assay in predicting cell type-specific binding sites across factors and conditions. In particular, given that the recent advances in genomic technologies continue to make chromatin accessibility interrogation more widely applicable, chromatin accessibility-based TFBS prediction methods will find more application in global-scale gene regulation analysis, such as cell type-specific CRM identification, multi-lineage regulatory landscape comparison and be used as structural priors in global gene regulatory network inference (87). Our method combined cell type-specific regulatory information in chromatin accessibility data with a range of genomic evidences in *cis* to drive more accurate binding site prediction. This lessened reliance on multiple expensive data-types in *trans* allows our method to be more readily scaled to a larger number of TFCT conditions.

In addition to building useful classifiers for global-scale TFBS mapping, our study aimed to identify factors that distinguish motif sites that are bound from the vast majority of unbound motif sites and provide mechanistic insights into TF binding dynamics and diversity. While chromatin accessibility assays allowed careful descriptions of the chromatin environment around TF binding sites, the sequence environment that fosters TF binding specificity and cooperativity, in contrast, is arguably harder to unravel and has thus remained by and large a conjecture that needs to be disentangled and tested in a more systematic fashion (91). Our trained sparse logistic regression models encapsulate some of the diverse combinations of sequence features that lead to TF-specific binding, such as GC/CpG content plays in binding site prediction. Additional features that describe motif-proximal sites, such as k-mer features and predictors of DNA shape, need to be investigated in more diverse biological contexts (71, 92, 93). For CpG features in particular, cell type-specific methylation assays, such as BS-sequencing could bring more functional relevance to its predictive modeling (48). Our study also provides a framework for examining co-clustered binding sites in a relatively unbiased manner; a key avenue of investigation, as binding site clustering is required for activation at many well studied loci (4, 49, 94, 95).

Evolutionary conservation represents yet another type of data that has long been associated with functional TF binding. We tested a range of conservation features in this work, including both measures of cross-species divergence and population-level polymorphism (e.g. SNP density) from ENCODE and 1000 genomes respectively (96). We found that SNP density at motif sites appear to have a diminished effect on binding site prediction (unpublished data), in comparison to cross-species divergence (97, 98). Among the cross-species conservation scores, mammalian conservation (both phyloP scores and phastCons scores) seemed the more relevant evolutionary distance than vertebrates or primates conservation (as evidenced by their selection in models for multiple TFs during MocapS training). This perhaps suggests a shift in balance between the regulatory conservation within mammalian species and the site divergence experienced among primates pointing to a precarious relationship between binding site conservation and divergence during evolution (55, 99).

As technical advances in genomics and statistics enable the accurate and large-scale mapping of a large number of TFBS, predictive methods that combine sequence features with chromatin accessibility modeling represent a promising direction for resolving the myriad binding sites across a diverse array of TFs and cell types. Large-scale mapping of TFBS, when connected with gene expression data, will in turn promote a better and more systematic understanding of mammalian gene regulation and enable large scale network inference via the generation of detailed structural priors (54, 88, 100).

## Supplementary methods

### Method Overview

Mocap combines information from motif, chromatin accessibility and a range of sequence-associated genomic features to predict TFBS. We start with a liberal set of candidate motif sites, and create a generic unsupervised method (MocapG) that ranks motif sites based on local accessibility. We then, building on this baseline method, adopt a supervised machine learning scheme (training sparse logistic regression classifiers MocapS) to investigate the contribution of different features to TFBS prediction and probe the diversity in TF binding behaviors. In training MocapS, we model both cell type-specific chromatin accessibility and footprint patterns around motif sites. We investigate (for different types of TFs) the contribution of sequence properties, such as GC/CpG content, surrounding motif sites to TFBS prediction. We use L1-regularized logistic regression to integrate motif-associated genomic features, which we find to be a good balance of performance and interpretability (in comparison to methods such as support vector machines, prior published methods, and methods adopting other regularization schemes). Lastly, we extend the trained classifiers of MocapS to TFCT conditions without ChIP-Seq label in a data-driven manner (MocapX). The goal is to create a TFBS prediction pipeline that balances method sensitivity with selectivity, and be able to generalize the specifically trained models to more factors and cell types via cross-sample TFBS predictions.

Below we detail the curation of motif models extracted from public databases, the computation of motif-associated genomic features, and the implementation of MocapS and MocapX.

### Motif models curation

We drew PWMs from two motif collections (Figure S1A). To reduce information redundancy for TFs with multiple motif representations, we filtered motifs based on pairwise PWM comparisons. Briefly, for a given motif pair, we vectorize the PWMs and calculate length-normalized Euclidean distance between the two vectors as follow

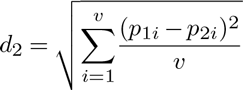

where *p*_1_ and *p*_2_ represent the length-normalized PWM vectors, *v* denotes the length of the vectorized PWMs. For motif pairs of unequal length, we let *v = min*(*p*_1_, *p*_2_) where we used the alignment that results in the smallest Euclidean distance among all possible ungapped PWM alignments. For motif pairs that are highly similar (with normalized Euclidean distance *<* 0.14), the PWM of lower entropy was retained (Figure S1B-D).

### Detecting motif-associated footprint patterns from chromatin accessibility data

To assess the probability that a footprint profile (*F*) exists around a motif site, we used a pair of binomial tests to gauge whether the motif region is protected from enzyme digestion as compared to its left and right flanking regions (4).

For DNase-Seq, we scored for the depletion of DNase I cuts at motif sites in a strand-specific manner, as strand imbalance is often conspicuous because of the size selection steps in DNase-Seq protocol (Figure S6). Specifically, on the positive strand, we assessed if a motif region is depleted of cuts as compared to the left flanking region.

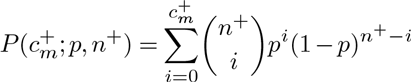

On the negative strand, we assessed if a motif region is depleted of cuts as compared to the right flanking region.

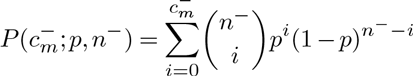

where 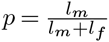. *p* denotes the expected size-normalized cut probability, where *l*_*m*_ and *l*_*f*_ denotes the size of the motif and its flanking region respectively. We empirically set the size of flanking region as 1.75 the size of motif length to optimize ChIP-Seq peak recovery. 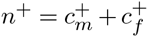 is the total number of cut counts within the motif site 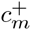 and its left flanking region 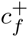 on the positive strand, whereas 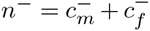 is the total number of cuts within the motif site 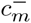 and its right flanking region 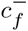 on the negative strand.

For ATAC-Seq, as the strand imbalance immediately flanking binding motif is less pronounced, we merged strand information to assess if a motif region is depleted of cuts as compared to the left and right flanking region respectively.

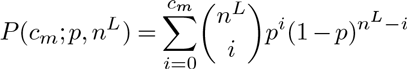

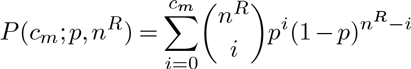

where 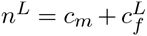 is the total number of cuts within the motif site *c*_*m*_ and its left flanking region 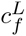 on both strands, whereas 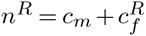 is the total number of cuts within the motif site *c*_*m*_ and its right flanking region 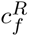 on both strands.

The footprint score was calculated for the DNase I footprint as

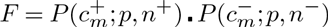

and for the ATAC-Seq footprint as

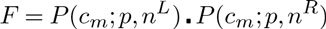

We shuffled the per base-pair cut count data around each site and reran the same binomial tests 500 times. A *p*-value threshold corresponding to a false discovery rate of 0.01 was chosen as the threshold for determining whether a footprint profile exists for each motif site.

### Profiling motif-associated sequence features

To profile the sequence environment surrounding TF binding motifs, we calculated GC content and CpG frequency for a region 100bp upstream and downstream of each motif site. We also computed a binary CpG island feature to assess if a motif resides in a larger region of inflated CpG count. A CpG island was assessed with the following criteria (2, 5)

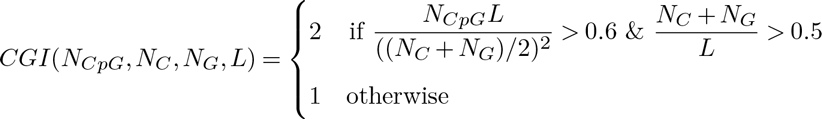

where *L* is the size of the region encompassing 400bp upstream and 400bp downstream of the length of a motif, *N*_*C*_, *N*_*G*_ and *N*_*CpG*_ are the total count of C, G and CpG dinucleotide respectively.

### Sparse logistic regression training (MocapS)

We trained sparse logistic regression models to classify candidate motif sites as true (motif overlapping with ChIP-Seq) or false (motif not overlapping with ChIP-Seq peaks) binding sites using a range of motif-associated genomic features, including motif matching quality, chromatin accessibility, TF footprint, TSS proximity, evolutionary conservation, motif region mapability and sequence-associated GC/CpG features. The goal is to build an ensemble of TFCT-specific and yet generalizable predictive models that address the differences in TF binding preferences and facilitate cross-sample binding site prediction. The logistic regression model is specified as below:

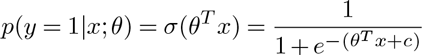

where *x* represents motif samples, *y* ∊ [1, −1] corresponds to bound and unbound motif labels respectively; *θ* specifies the 30 motif-associated genomic features that are mostly highly correlated with the ChIP-Seq feature in the tested motif set and cell type environment (after two rounds of correlation filtering); *c* specifies the intercept. Features, such as SNP density, TES proximity and exon, have relatively low correlation with ChIP-Seq signals, and a weak contribution to the overall prediction of TFBS, so we excluded them from the model *a priori* for all TFCTs.

To create generalizable models and ensure only a most relevant subset of the genomic features are included in the logistic regression model for each TFCT, we specified a sparsity constraint to minimize the below objective function with L1 regularization

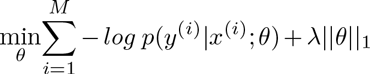

where *M* is the number of candidate motif sites used for model training; λ is the shrinkage parameter corresponding to the sparsity of the trained logistic regression model where with L1-regularization, all but the most relevant features are shrink to zero. As the number of true binding sites takes up only 0.01%-5% of the total number of candidate sites (imbalanced class labels), we weighed the true and false motif classes to create a weight-balanced training dataset. For each TFCT condition, we tuned the shrinkage parameter λ to select a λ that optimizes the AUPR to achieve more sensible site ranking (also, AUPR scores are less biased than AUROC scores in assessing classification performance in imbalanced datasets). To avoid overfitting and promote model generalization, λ parameter was further tuned such that the resulting classifier is sparser than an optimally performing classifier with near-optimum (within one standard error of the optimum) cross-validation performance. We adopted a final bootstrap aggregation step to reduce estimator bias and help achieve more robust classifications.

### Implementation of cross-sample TFBS prediction (MocapX)

To extend the utility of the above sparse logistic regression classifiers ensemble to TFCT conditions where ChIP-Seq data is missing, we used robustly weighted least squares regression to derive a mapping vector *β* to assign new TFCT prediction problems to trained classifiers based on genomic feature distance. When minimizing the below error function

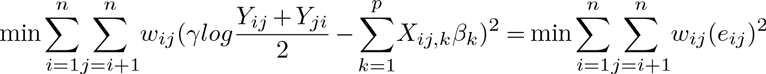

where *n* is the number of TFCT conditions; *p* is the number of features; *X*_*ij,k*_ is the binary probability that feature *k* from TFCT *i* and TFCT *j* are derived from the same distribution based on KS test for continuous features and Chi-square test for categorical features; *Y*_*ij*_ is the predictive performance (AUPR) of model trained in TFCT *i* when applied to classify motifs in TFCT *j.* We used predictive performance of MocapG as a baseline and used adaptive grid search to assign γ to seperate sample pairs that cross-predict well from those that predicts poorly. For DNase-Seq sample mappings, for example, we set γ as follows

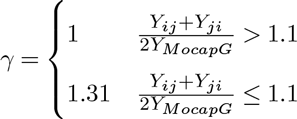

γ was re-optimized with the exclusion of footprint features, when deriving mappings between ATAC-Seq samples and DNase-Seq data trained models. *w* was fitted via iteratively re-weighted least squares regression with the Tukey’s bisquare family psi functions as below (1, 3).

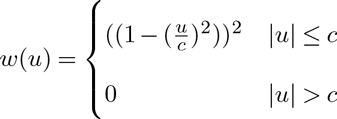

where *u* is the regression residual.

**Figure 1:**
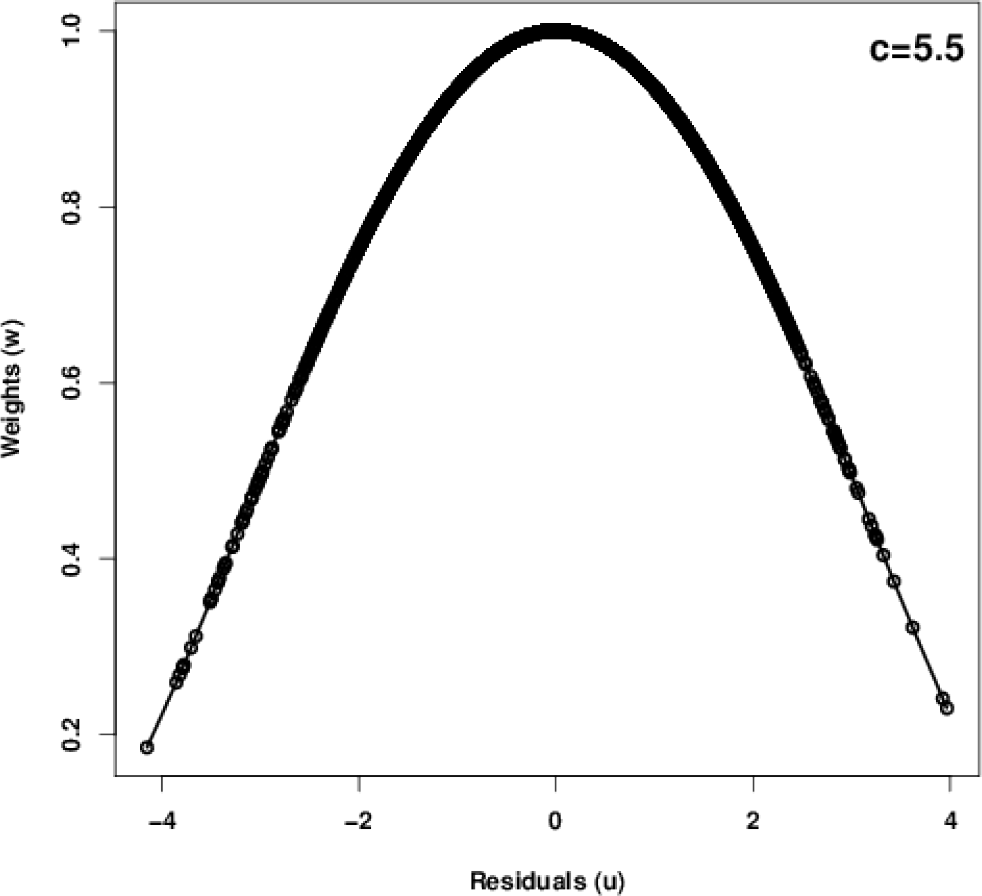
Tukey’s bisquare function specifies sample weights based on regression residuals.

We selected the bisquare psi function hyperparameter *c* based on cross-validation sample-mapping performance. Distances between feature vectors were then used to assign weight *w* to a hold-out TFCT condition based on the weights of its nearest neighbors. We used 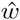 and 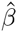 to compute a sample mapping *i* → *j* that maximizes 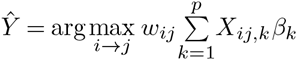, but to avoid mappings that do not significantly improve performance as compared to the generic method MocapG, we assign the mapping if and only if the estimated cross-prediction performance 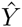 compares favorably with *Y*_*M ocapG*_ in TFCT *i* as below

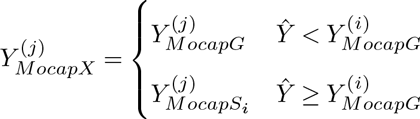

